# RNA plasticity emerges as an evolutionary response to fluctuating environments

**DOI:** 10.1101/2024.10.02.614758

**Authors:** Paula García-Galindo, Sebastian E. Ahnert

## Abstract

Phenotypic plasticity refers to the ability of a single genotype to produce multiple distinct phenotypes. Using the computationally tractable genotype-phenotype (GP) map of RNA secondary structures, we model RNA phenotypic plasticity using the Boltzmann distribution of secondary structures for each genotype. Through evolutionary simulations that involve periodic environmental switching on the GP map, we reveal that RNA phenotypes can adapt to these fluctuations towards an optimal plasticity. The optimal phenotypes exhibit dominant near-equal Boltzmann probabilities of distinct structures, each representing the fittest structure for each alternating environment. Our findings demonstrate that phenotypic plasticity, a widespread biological phenomenon, is a fundamental evolutionary response to changing environments for RNA secondary structure. We also find naturally evolved functional RNAs that exhibit optimal plasticity unlikely to arise by neutral drift alone, suggesting functional relevance in fluctuating environments.

**Author summary:** Phenotypic plasticity—the ability of an organism to change its traits in response to environmental variation—is a common concept in biology, but remains challenging to study quantitatively. In this work, we explore plasticity at the molecular level in RNA, a molecule essential both for storing genetic information and for performing cellular functions through its structure and sequence. We focus on how the structural plasticity of RNA can evolve in response to fluctuating environmental conditions. Using computational models, we show that RNA can evolve towards an optimal distribution of structures when exposed to environments that alternate over time. We show that this near-optimal plasticity is present in functional non-coding RNAs that have evolved naturally. We identify key factors, such as the rate of environmental switching and structural properties, that influence how RNA plasticity emerges and evolves. Our findings offer new insights into molecular-level plasticity and suggest that RNA’s structural adaptability may contribute to its biological function.

## Introduction

From global climatic changes to local fluctuations of temperature or chemical conditions, environmental shifts have continuously driven biological organisms to evolve strategies to cope with the challenges of a varying environment [1]. A substantial body of theoretical research on evolution in changing environments originates with Levins’ foundational work [2], which first introduced the concept of the “fitness set” and more broadly explored how environmental fluctuations can hinder optimal adaptation to a single environment. Others have examined the impact of changing environments on performance trade-offs, using the Pareto front to describe optimal phenotypic solutions [3–5]. Further theoretical models have investigated the evolution of reaction norms, which describe a varying phenotypic response as a function of the environment [6, 7]. Such a varying phenotypic response that arises from the same genotype is often referred to as phenotypic plasticity, and is a widespread phenomenon in biology [1]. However, it is still difficult to study the evolution of plasticity using quantitative models relevant to biology [8]. Here, we study the evolution of plasticity in a changing environment using biologically realistic phenotypes in the form of RNA secondary structures through the genotype-phenotype (GP) map framework. The RNA GP map is often used to systematically quantify general properties of evolutionary spaces, offering generality [9], while retaining biological relevance since the results apply to natural functional RNAs [10].

The GP map of RNA secondary structure has been studied extensively [11–15] and its large-scale structural properties have been characterised [16]. The nucleotide sequence represents the genotype, which is subject to mutations, while the RNA secondary structure represents the phenotype. This GP map can be computationally constructed for short RNA (*L <* 100) to limit the number of sequences (4*^L^*) involved and to approximate the three-dimensional RNA fold by its base pairings, or secondary structure, which can be easily computed through experimentally validated folding algorithms [17], and represented in dot-bracket notation [16]. The RNA secondary structure GP map is computationally tractable while still being biologically relevant, as the biological functions that short RNA performs in the cell (e.g., tRNAs, miRNAs) are often highly dependent on the folded structure [18, 19].

The large-scale structural properties of the RNA GP map appear to hold across several other GP maps too, including the HP model of protein folding, gene regulatory networks (GRNs), and other biological systems. These properties have a direct impact on evolutionary trajectories, as the accessibility of phenotypes via mutations is strongly affected by the shape and size of neutral networks in the GP map [9, 14–16]. Additionally, the structure of GP maps results in highly navigable fitness landscapes, implying that true fitness valleys are uncommon in nature [20]. The structure of GP maps has also been linked to the ubiquity of symmetry in structural phenotypes [21].

Most of the literature on the RNA GP map only considers one phenotype per genotype, the most stable or minimum free energy (MFE) secondary structure, even though RNA has inherent phenotypic plasticity. At constant temperature, the folding process of an RNA nucleotide strand is probabilistic due to thermal fluctuations at the molecular scale, allowing the RNA strand to dynamically fold and unfold over time, switching between different structures based on their respective free energies [22]. This ability of the same sequence to adopt different structures can be crucial for RNA’s cellular functions, such as its involvement in gene expression [23, 24], and a recently proposed ‘Non-deterministic GP map’ (ND GP map) framework aims to explicitly describe GP maps in which each genotype is mapped to a probability distribution of biological outcomes [25].

Some RNA sequences exhibit a high degree of plasticity, which means that the phenotype ensemble of probable structures consists of several low-energy folds separated by small energy gaps. Other sequences are associated with low plasticity, which corresponds to a large energy gap above the MFE structure. In previous studies on RNA secondary structure ensembles, the mutational robustness of an MFE structure has been shown to correlate with its structural similarity to other structures within the phenotype ensemble [26]. Furthermore, there is a correlation between an MFE structure and the suboptimal structures in the phenotype ensembles of one-point mutants, a relationship known as plastogenetic congruence [22]. These correlations help explain the reduction in plasticity observed for RNA in evolutionary simulations under constant environments [22]. This outcome is intuitive: if a biological function depends on a specific RNA structure being thermodynamically stable, it must also evolve to maintain mutational stability.

However, in fluctuating environments, RNA gentoypes may benefit from the ability to display more than one phenotype, in other words, whether it might become beneficial for plasticity to evolve. To investigate this phenomenon under simple assumptions, we measure the degree of plasticity that evolves in an evolutionary model of RNA secondary structures with two periodically switching environments, using the ND GP map for RNA of length *L* = 12 [25], which includes probabilistic folding in a many-to-many map from sequences to the Boltzmann ensembles of folds. We find that plasticity does indeed evolve in rapidly switching environments, as plasticity increases towards an optimal solution that narrows the plastic repertoire to a balanced distribution, representing equal probabilities of the two target RNA folds (see Fig.1). We show furthermore how the evolution of plasticity depends on the timescale of environmental change, the sampling frequency of the phenotypic distribution, and the generational timescale and mutation rate of evolution, as well as properties of the time-varying evolutionary target structures. Finally, we analyse the plasticity of functional RNA in the fRNA database, in which we find a number of short fRNAs approaching optimal plasticity.

**Fig 1.**
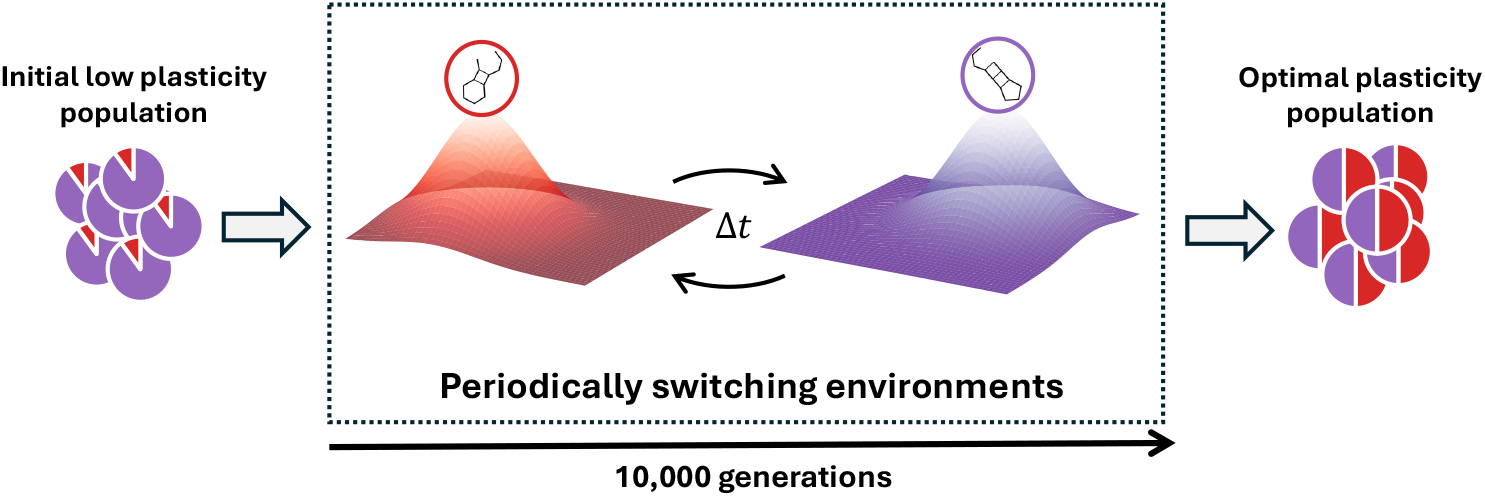
Outline of the evolutionary RNA model. Individual pie charts represent genotypes and the associated relative Boltzmann probabilities of the two target phenotypes, *T*_0_ (red) and *T*_1_ (purple). The initial population (left) undergoes evolution for 10,000 generations under periodically changing environmental conditions, implemented by two alternating fitness landscapes in which the respective fitness peaks correspond to the two target structures *T*_0_ and *T*_1_. The switch occurs every Δ*t* generations. We investigate whether this process results in genotype plasticity, in the form of an equal representation of the target structures in the Boltzmann ensemble.

## Methods

### The RNA non-deterministic genotype-phenotype map

The RNA ND GP map maps all possible RNA sequences of a fixed length to their respective Boltzmann ensembles of secondary structures represented in dot-bracket notation, and connects each sequence to its one-mutation neighbours to form a graph in which sequences are the nodes [25]. The phenotype associated to each genotype will be the Boltzmann ensemble found for a certain energy range (so that as Δ*E →* 0 only the MFE remains), and the Boltzmann probabilities are normalised for that set of structures [25]. Therefore, each sequence *g* folds to a secondary structure *p* with a certain Boltzmann probability *P* (*p|g*) = exp((*G_ens_ − G_p_*)*/k_B_T*), where *G_p_*is the free energy that corresponds to structure *p*, and *G_ens_* is the ensemble free energy: *G_ens_* = *−k_B_T ·* ln*Z* with partition function *Z* = Σ_p_ exp(*−E_p_/k_B_T*), where *E_p_* is the specific energy associated with structure *p* [22].

In other words, at any point in time in an RNA population of genotype *g*, the number of RNAs that fold to a certain phenotype *p* will be proportional to the Boltzmann probability *P* (*p|g*). In this study, we fix the RNA length to *L* = 12, so we construct the RNA12 ND GP map. For each RNA nucleotide sequence, the secondary structure ensemble, its ensemble free energy, and the free energy of each structure within the ensemble are computed using the `ViennaRNA subopt` [17] function set to default parameters (e.g. temperature *T* = 37*^◦^*C, no isolated basepairs) and energy range above the minimum free energy as Δ*E* = 15*k_B_T*, like in previous works [25].

### Evolutionary dynamics

#### Target secondary structures

To select target secondary structure pairs {*T*_0_*, T*_1_} for periodic switching during an evolutionary simulation, we categorise the pairs based on the phenotypic frequencies or neutral space sizes *NSS* of the respective targets in the RNA12 ND GP map, as well as their structural difference measured in terms of their Hamming distance to each other.

First we categorise all the possible secondary structures of the RNA12 ND GP map in terms of their *NSS*: large (*∼* 10^5^) and small (*∼* 10^3^). We then compute all the possible target pairs that result from pairing secondary structures within and between these two *NSS* size categories, and therefore bin the target pairs into three *NSS* pair categories: both large, mixed (i.e. one large, one small), and both small. For each target pair in each category, the Hamming distance between targets is calculated, where the Hamming distance between two secondary structures of length *L* is calculated using 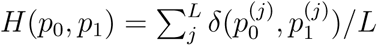, with *p*^(^*^j^*^)^ being a nucleotide base in dot-bracket notation and in our case *L* = 12. We then bin the pairs in each *NSS* category by their Hamming distance *H* (large, medium and small). For our RNA12 data, to make sure there are enough pairs distributed in each Hamming distance category, we choose the *H* bins to be the following: large *H ∈* [0.58, 1.0], medium *H ∈* [0.33, 0.58], and small *H ∈* [0, 0.33].

The target secondary structure pairs in these categories have generally a small overlap in genotype space on the ND GP map (see Fig. 2). In this case, our target pairs share less than 1% of genotypes, which means that plasticity solutions are overall rare, implying that it can be dependent on the evolutionary dynamics and ND GP map structure whether these genotypes are found. There is a moderate correlation of *r* = 0.46 between the genotype overlap size and total genotype size for a target pair.

**Fig 2.**
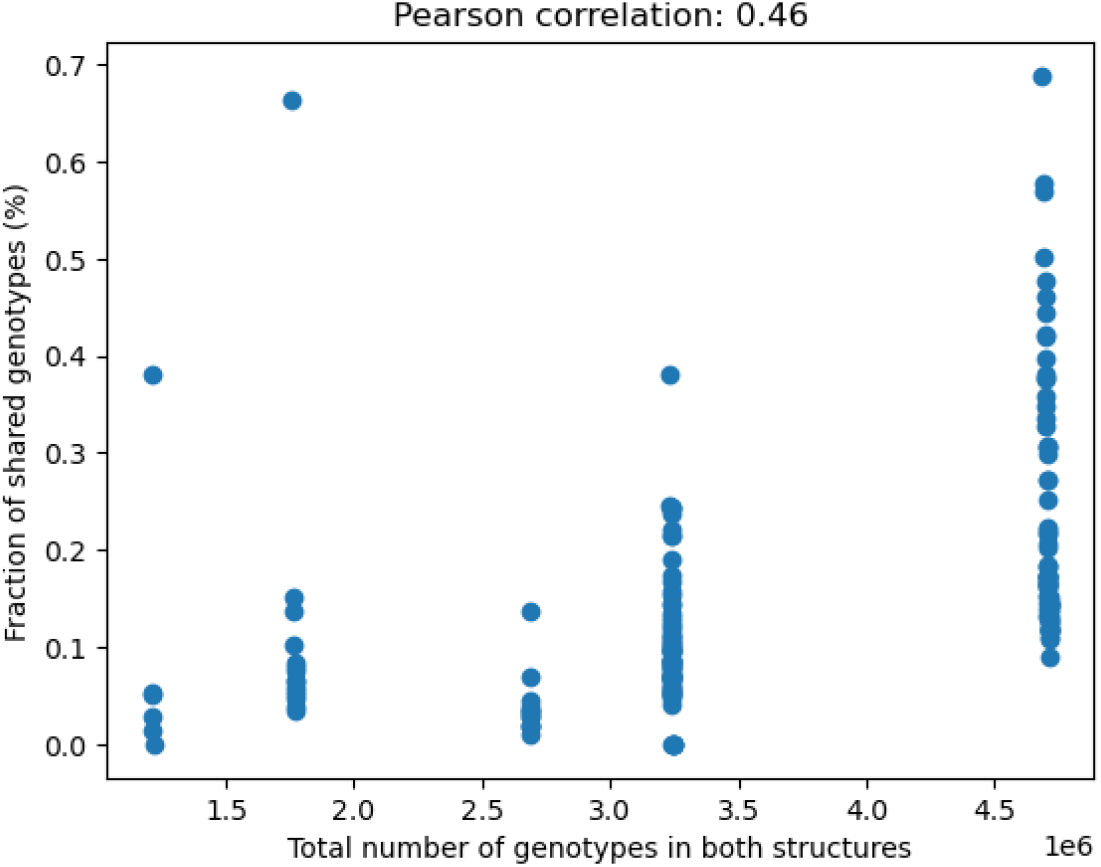
Plastic genotypes are rare. Less than 1% of genotypes from two structures in a target pair have plasticity, and this has a moderate correlation, Pearson *r* = 0.46, with the size of genotype space that the two structures occupy.

#### Genotype populations

Using a specified target secondary structure pair {*T*_0_*, T*_1_*}* we define *plastic* genotypes, *optimal plasticity* genotypes, and *low plasticity* genotypes. A plastic genotype is one that contains both targets in its Boltzmann ensemble. An optimal plastic genotype is one with target Boltzmann probabilities (*P*_0_*, P*_1_) = (0.5, 0.5), which represents the midpoint of the Pareto front connecting the archetypes, *P*_0_*_,max_* = (1.0, 0.0) and *P*_1_*_,max_* = (0.0, 1.0) [4]. A low plasticity genotype of either target is one that maximises the probability of adopting that target structure among all the plastic genotypes.

#### Model

Starting with a specified target pair {*T*_0_*, T*_1_*}* and a homogeneous low plasticity genotype population of size *N* = 100 (such that *Nµ* spans polymorphic, *≫* 1, and monomorphic, *≪* 1, regimes), we navigate the sequence space of the RNA12 ND GP map over time through mutations and sampling from the Boltzmann ensembles of mutated sequences to generate phenotypes that then undergo a selection process. The evolutionary dynamics follow the classical haploid Wright-Fisher model [27], where in each generation a fixed-size genotype population undergoes point mutations with a mutation rate *µ* (probability of a mutation happening per genotype base at each generation step) and genotypes are selected with replacement using roulette-wheel selection, where the probability of being chosen is proportional to their fitness. To evolve RNA in fluctuating environments we periodically switch between the fitness landscapes of the two target structures with a switching frequency of 1*/*Δ*t*, meaning a switch occurs after a time interval Δ*t*.

During each target epoch, the target secondary structure is assigned a maximum fitness value of *f* = 1.0, while the fitness of other secondary structures is determined by the Hamming fitness *f_H_i__* = 1 *− H*(*p_i_, p_T_*), where *p_i_* is the secondary structure for which fitness is being calculated, and *p_T_*is the target secondary structure, both represented in dot-bracket notation. A fitness landscape is constructed by assigning Hamming fitness values to all phenotypes in the RNA12 ND GP map relative to a specific target structure. For each genotype, a set of *N_p_*secondary structures is sampled from its Boltzmann ensemble, with each fold assigned a corresponding Hamming fitness value. The genotype’s fitness is then defined as the maximum fitness among the sampled phenotypes, effectively giving the genotype *N_p_* opportunities to realise a high-fitness conformation within a given time frame. This maximum-fitness sampling approach reflects a biologically motivated assumption. RNA molecules dynamically sample an ensemble of structures due to thermal fluctuations, and interactions with molecular partners often depend on the adoption of specific conformations [28]. For example, if an RNA’s function is to bind a ligand or interact with a protein, it is the conformations that are most competent for this interaction that confer a selective advantage. In this sense, selection acts on functional episodes rather than on a time-averaged structure. The use of the maximum fitness among sampled phenotypes therefore serves as a proxy for this behaviour, capturing the idea that even transient access to a highly functional conformation can be sufficient to contribute to fitness.

Simulations run for *t* = 10, 000 generations, and all evolutionary trajectories are averages computed from sample sizes of *N_s_*= 100 runs. The evolutionary dynamics simulation is summarized in the following steps:

**0.0** Fix parameters *µ*, Δ*t* and *N_p_*.
**0.1** Generate initial low plasticity population that maximises *P*_1_ (*P*_0_).
**0.2** Set target to *T*_0_ (*T*_1_).
**1.1** For each genotype in the population, sample a number *N_p_* of folds from the phenotype Boltzmann ensemble determined by the RNA12 ND GP map.
**1.2** Assign Hamming fitness, *f_H_i__* = 1 *− H*(*p_i_, p_T_*), with respect to the target fold *p_T_* to each sampled secondary structure.
**1.3** Assign fitness of each genotype *f_i_* to be the maximum fitness in its *N_p_* set.
**2.** Generate a new population by sampling with replacement *N* genotypes with probability proportional to the fitness, 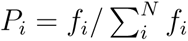
**3.** For each base position *j* in each genotype *i*, apply a random mutation with a Bernoulli probability *µ* (the point mutation rate). When a mutation occurs at 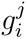, a random base different from the current one is chosen from {*A, C, G, U*} with uniform probability.
**4.** Increase the generation time by one step, *t → t* + 1, and set the mutated population as the updated population for *t*.
**5.** Repeat steps **1**-**4**, and at each Δ*t* generation steps implement a switch to the alternative target *T*_0_ *↔ T*_1_.
**6.** End simulation at *t* = 10, 000.

### Pareto front of RNA suboptimal structures

In biological systems, phenotypes optimize their performance for specific tasks through evolutionary processes. But when a phenotype needs to perform not only one task but multiple, the Pareto front framework can be translated to evolutionary trade-offs, such that its performance will evolve to allow for all tasks but cannot be the best at a single one [4]. To determine the resulting phenotype, a geometric framework was shown such that best–trade-off phenotypes are weighted averages of archetypes (phenotypes specialized for single tasks in question).

To model Pareto fronts in phenotype space, we consider a system performing multiple tasks, where each task *i* is associated with a performance function *P_i_*(*v*) defined over a continuous phenotypic space. We assume the following:

1. Single optimal phenotype per task: For each task *i*, there exists a unique phenotype 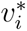 (the archetype) that maximizes performance in that task.
2. Performance declines with distance from the archetype: Performance for task *i* decreases as the phenotype deviates from 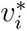. Distance is defined as a metric based on an inner product norm:

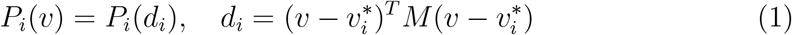

where *d_i_* represents a distance metric between phenotype *v* and archetype 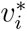, and *M* is a positive-definite matrix defining the norm. When *M* = *I*, *d_i_* reduces to the standard (squared) Euclidean distance: 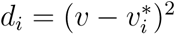.

Under these assumptions, the geometry of the Pareto front emerges naturally. Generally, for *N* tasks, the Pareto front forms a convex polytope (a simplex) whose vertices are given by the archetypes 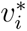 [4]. Phenotypes located on this front represent optimal trade-offs between tasks, where improving performance in one task necessarily results in performance loss in at least one other task. This geometric framework links evolutionary trade-offs directly to the structure of phenotype space, providing a quantitative description of specialization and generalism in multi-task systems [4]. In particular, for two tasks (*M* = 2), the Pareto front corresponds to the straight line segment connecting the two archetypes. The optimal phenotype will be on this segment, and the exact position on this line relates to how important the two tasks are to the evolutionary process: the closer to an archetype, the more important that task was for survival.

We use this framework to define a Pareto front with respect to the probabilities of the target structures in the Boltzmann ensemble of an RNA sequence. A Boltzmann ensemble with target structure of probability 1 is defined as an archetype of the respective target, a theoretical point because it is not thermodynamically feasible to have only one structure at *T* = 37*^◦^* and energy range Δ*E* = 15*k_B_T*. In our alternating environments model, since the time interval Δ*t* is fixed, we assume equal contribution between the two archetypes for survival, such that the Pareto front optimal plasticity point becomes the point that balances both probabilities, (*P*_0_*, P*_1_) = (0.5, 0.5).

### Functional RNA

The functional RNAs are found in the database fRNAdb [29]. We choose to analyse the fRNAs that are of length *L ≤* 50, where the total number of fRNAs in this length range is 237,461. The site-scanning method [30] is used to create a sample of 5,000 plastic mutational neighbours for each analysed fRNA sequence:

**1.** For each analyzed sequence, store the MFE structure and the first suboptimal structure as *T*_0_ and *T*_1_, and their Boltzmann probabilities *P*_0_ and *P*_1_.
**2.** Start from the first base in the sequence and proceed from left to right.
**3.** Mutate the selected base by randomly introducing a mutation (i.e., substituting it with a different nucleotide: A, U, G, or C).
**4.** Check whether the mutated sequence is a plastic gentoype, i.e. its plastic repertoire still includes both *T*_0_ and *T*_1_.

**4.1** If both *T*_0_ and *T*_1_ are present, record their Boltzmann probabilities 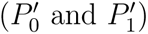 for the mutated sequence.
**4.2** Update the sequence being analyzed to the mutated sequence, and proceed to analyse the next nucleotide position with **3-4**.
**5.** If either *T*_0_ or *T*_1_ is missing from the repertoire, do **3-4.** until all different mutations of that nucleotide are checked.
**6.** If none of the mutations give a plastic genotype, move to the next base position in the sequence and repeat the procedure **3-4**.
**7.** End when 5,000 plastic genotypes are recorded (see Supplementary Information S1).

Next, for each sequence, we use the site-scanning values 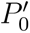 and 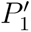 to generate a probability distribution of distances to optimal plasticity, denoted by *d*:

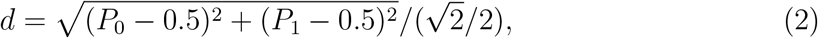

which measures the Euclidean distance of the observed plasticity pair (*P*_0_,*P*_1_) from the optimal plasticity point (0.5,0.5), normalized to lie in the interval [0, 1].

We choose to analyse for significance the fRNAs that are highly plastic, up to *d* = 0.35. Then we calculate an empirical p-value for each sequence by comparing its distance *d* against the distribution of distances *d^′^* from the 5000 site-scanning neighbours:

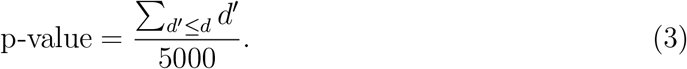

A p-value less than 0.05 indicates that the fRNA’s plasticity is significantly close to optimal, and unlikely to arise by chance under mutation.

Since we are computing significance tests for all the fRNAs with distance *d* up to *d* = 0.35, we correct for multiple hypothesis testing by implementing a Benjamini-Hochberg correction at 10% [31]. The total sequences up to the cutoff *d* = 0.35 are 3, 685, while the number of significant sequences we find is 2, 412.

## Results

### Optimal plasticity evolves in switching environments

#### Rapid environmental switching promotes optimal plasticity

To begin investigating the evolution of plasticity we illustrate the prevalence of plastic phenotypes across a wide range of values for mutation rate *µ* and switching time Δ*t*, through an example target pair 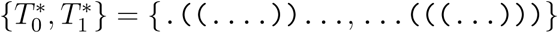, where both targets are relatively different (large Hamming distance between them, *H* = 0.83) and both are common in the GP map (large neutral space sizes), see Methods. The initial low plasticity population of size *N* = 100 was chosen such that *P*_1_ is maximised: AUCGGCCGGGCC with {*P*_0_, *P*_1_} = {0.001, 0.893}, and the fitness sampling number is *N_p_* = 10.

The time series in Fig.3 shows the evolution of abundances of the different genotypes (c.f. Methods) and mean fitness for *t* = 10, 000 generations for different values of the mutation rate *µ* and switching time Δ*t*. The abundances correspond to the plastic genotypes, *x_p_* (orange), with both targets in their phenotype ensembles; the genotypes with only the targets *T*_0_ or *T*_1_ in their phenotype ensembles, *x_T_*_0_ (red) and *x_T_*_1_ (purple); and finally the other possible genotypes *x*_0_ (brown), which are treated as noise. There is a noticeable increase in plastic genotypes and fitness at high switching frequency (low Δ*t*). In Fig. 4, two examples are picked from Fig. to show the impact of two different switching frequencies on the initial behaviour of the plastic genotypes’ abundance. Both cases have mutation rate *µ* = 0.001, while (i) has Δ*t* = 25 and (ii) has Δ*t* = 100. The simulation starts at *T*_0_ region, so we expect the *T*_0_ population abundance to increase with the mean fitness. However, in (i) of low Δ*t* the population of *x_p_* quickly adapts.

**Fig 3.**
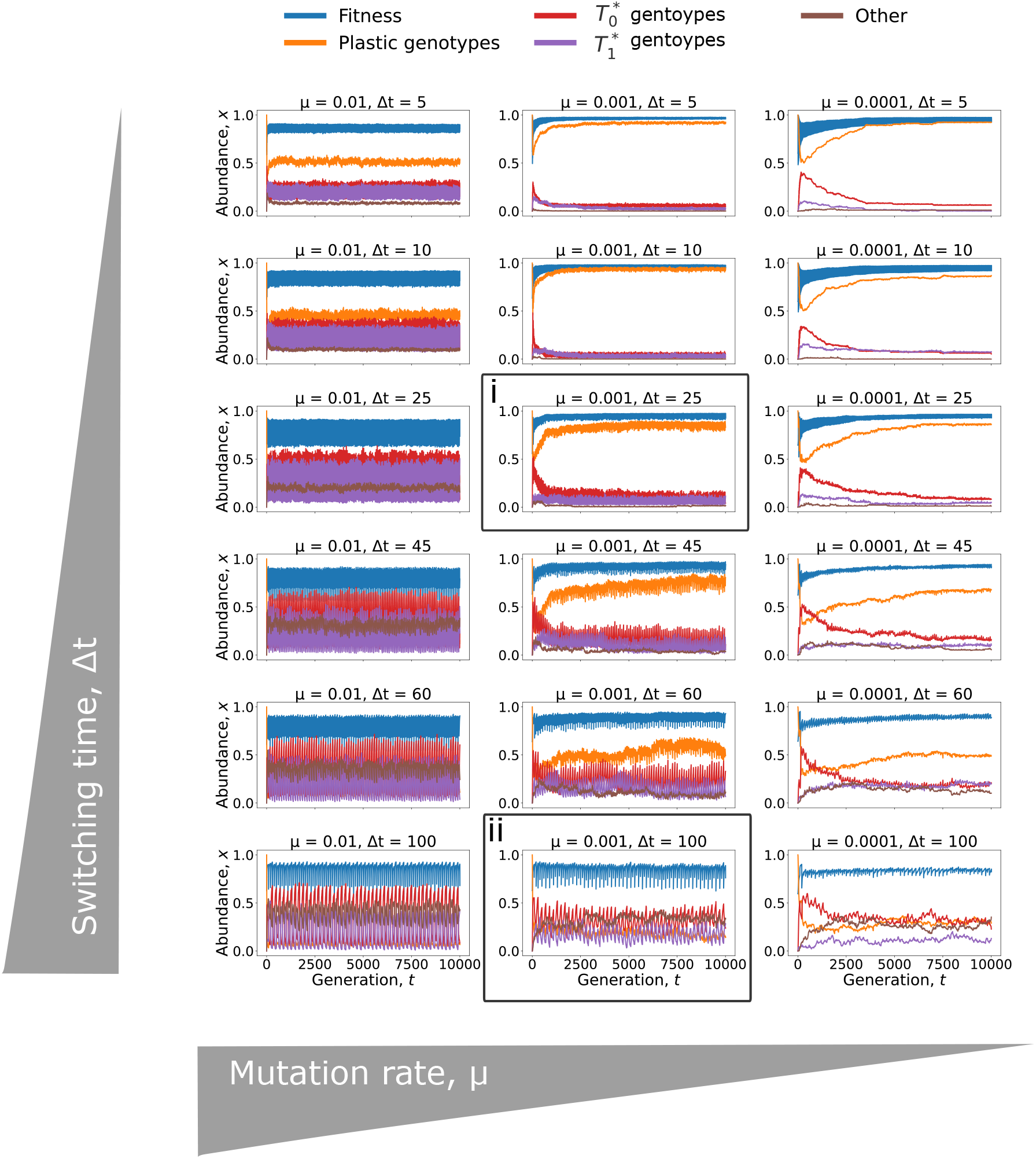
Example showing evolution of plastic genotypes depending on the mutation rate and switching time in periodically switching environments. Time series simulations are shown for target pair 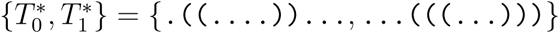. These targets have a large Hamming distance, *H* = 0.83, and large neutral space sizes, ranked 12th and 6th, respectively, out of the total 271 in the RNA12 ND GP map [25]. The initial population maximises *P*_1_: AUCGGCCGGGCC with *P*_0_*, P*_1_ = 0.001, 0.893. The time series are shown for different mutation rates *µ* and Δ*t* switching times. The *x*-axis of each subplot is time *t* generations up to 10, 000 and the *y*-axis is the abundance *x* of the different types of genotypes and the mean fitness value. Genotypes are divided into plastic genotypes with both targets in Boltzmann ensemble, the genotypes with only *T*_0_ or only *T*_1_ in their ensembles, and the other genotypes are treated as noise (see Methods).

**Fig 4.**
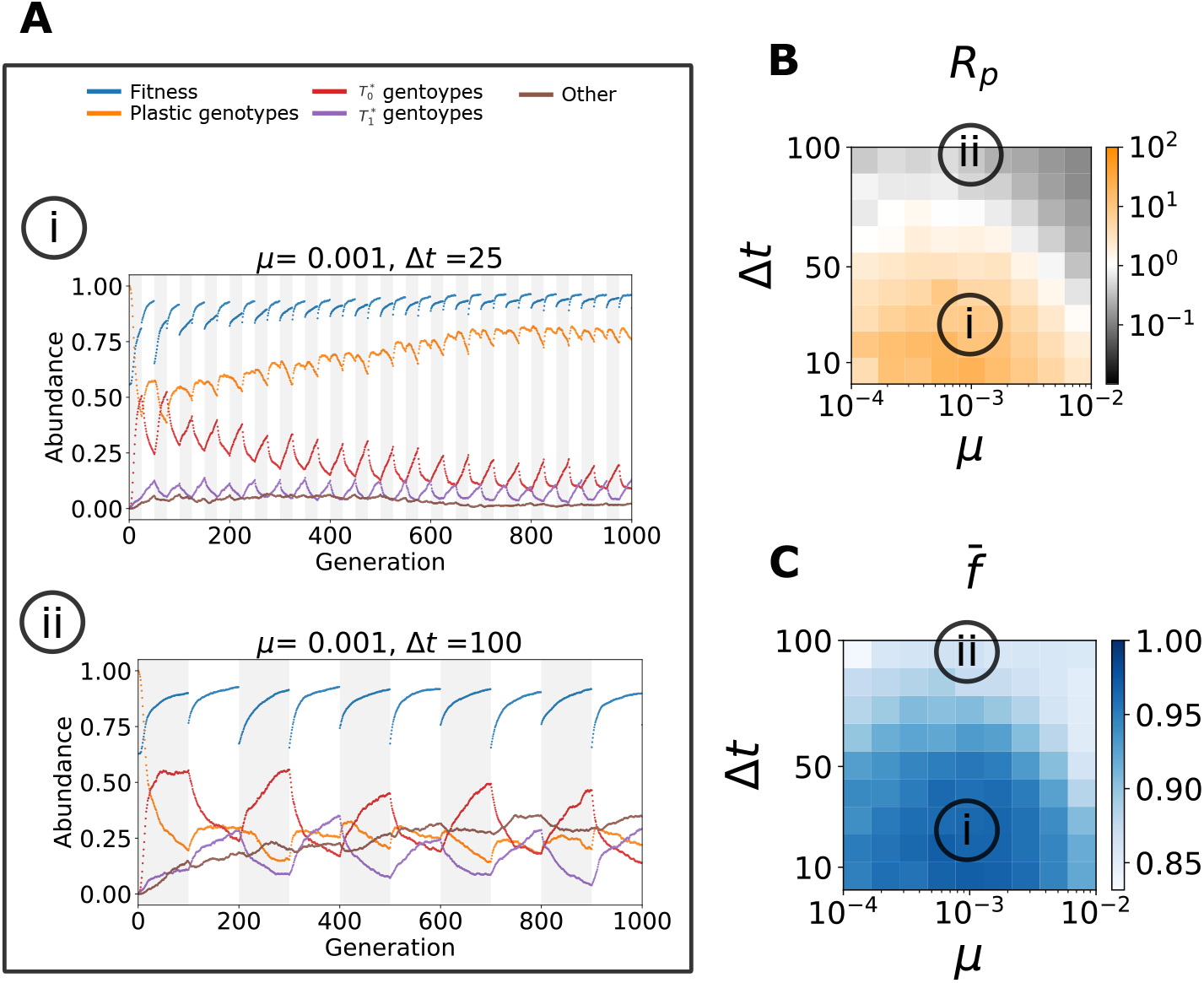
Enlarged examples from Fig. showing that plastic genotypes emerge in rapidly switching environments as an adaptive response. **(A)** Enlarged versions of two plots in Fig. 3, marked (i) and (ii) for the time range *t* = 0 to *t* = 1, 000. The grey (white) region highlights the switching time corresponding to each target *T*_0_ (*T*_1_). The regions switch every Δ*t*. **(B)** Parameter sweep heat maps for different (*µ,* Δ*t*) of the plastic abundance ratio *R_p_*and **(C)** of the mean fitness *f̅*. These quantities are time averages from the midpoint to end of the simulation. The heat maps have a strong positive correlation of Pearson *r* = 0.75. The map for *R_p_* shows that plasticity is prevalent (Plastic abundance ratio *>* 1 = orange shades) for a wide range of mutation rates *µ* and relatively fast switching, Δ*t <* 50. The specific simulations (i) and (ii) of (*µ,* Δ*t*) = (0.001, 25) and (*µ,* Δ*t*) = (0.001, 100) respectively are highlighted on the maps.

To validate that generally a high switching frequency makes the plastic genotype population adapt, we compute the plastic abundance ratio: *R_p_*= *x_p_/*(*x_T_*_0_ + *x_T_*_1_ + *x*_0_), which is a ratio of the accumulated abundance from the midpoint of the simulation until the end, of the plastic abundance *x_p_* over the rest of genotypes. We assume the midpoint as a reasonable time proxy at which the time series stabilises, thereby avoiding the computational expense of computing the exact stationary state for each simulation, while being consistent throughout the analysis.

We define the plastic population as adapted if *x_p_ >* 0.5, so if *R_p_ >* 1 the plastic genotypes have adapted, but if *R_p_ <* 1 the rest of the genotypes dominate. For the previous example, we show different *R_p_*(*µ,* Δ*t*) in Fig.4B, in which we observe that for a fixed *µ*, decreasing Δ*t* leads to an increase in *R_p_*, in other words faster switching between environments increases the prevalence of plasticity. In addition, Fig. 4C shows a positive correlation (Pearson *r* = 0.75) between *R_p_* and mean fitness *f̅*, verifying that the plastic population allows quick access to the target secondary structure, and therefore is adaptive.

After having established the prevalence of plasticity under certain conditions we now investigate the degree of plasticity, by analysing the population averages of the target Boltzmann probabilities *P̅*_0_ and *P̅*_1_. We show the evolution for examples (i) and (ii) in Fig.5A, where the Pareto front [4] in the phase diagram of (*P̅*_0_,*P̅*_1_) is the line connecting the archetypes *P̅*_0_*_,max_* = (1.0, 0.0) and *P̅*_1_*_,max_* = (0.0, 1.0), points that represent maximum fitness in each respective target fitness landscape. Since both targets equally contribute to the evolutionary process, the optimal phenotype on the Pareto front is located at the midpoint of the line connecting the two archetypes [4]. To verify that high switching frequency leads the genotypes towards optimal plasticity, we compute the normalised Euclidean distance 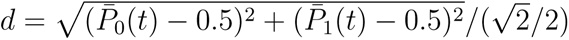, from the last point (*P̅*_0_(*t*)*, P̅*_1_(*t*)) where *t* = 10, 000, to the midpoint (0.5, 0.5). As expected, both examples (i) and (ii) get closer to the optimal point, and example (i) is closer to the midpoint than example (ii). The distribution of target probabilities at *t* = 10, 000 of the genotype population is shown for all *N_s_*= 100 simulations, verifying that probabilities in (i) are localised closer to the midpoint, while the ones in (ii) are localised on the axes. Finally, we verify that the conditions that maximise plasticity (minimise *d*) correlate with maximising the plastic population abundance and the mean fitness: the parameter sweep of the time averaged distance *d̅* is shown in Fig. 5B, and has Pearson *r* = *−*0.63 with *R_p_* and Pearson *r* = *−*0.86 with *f̅* in Fig.4C.

**Fig 5.**
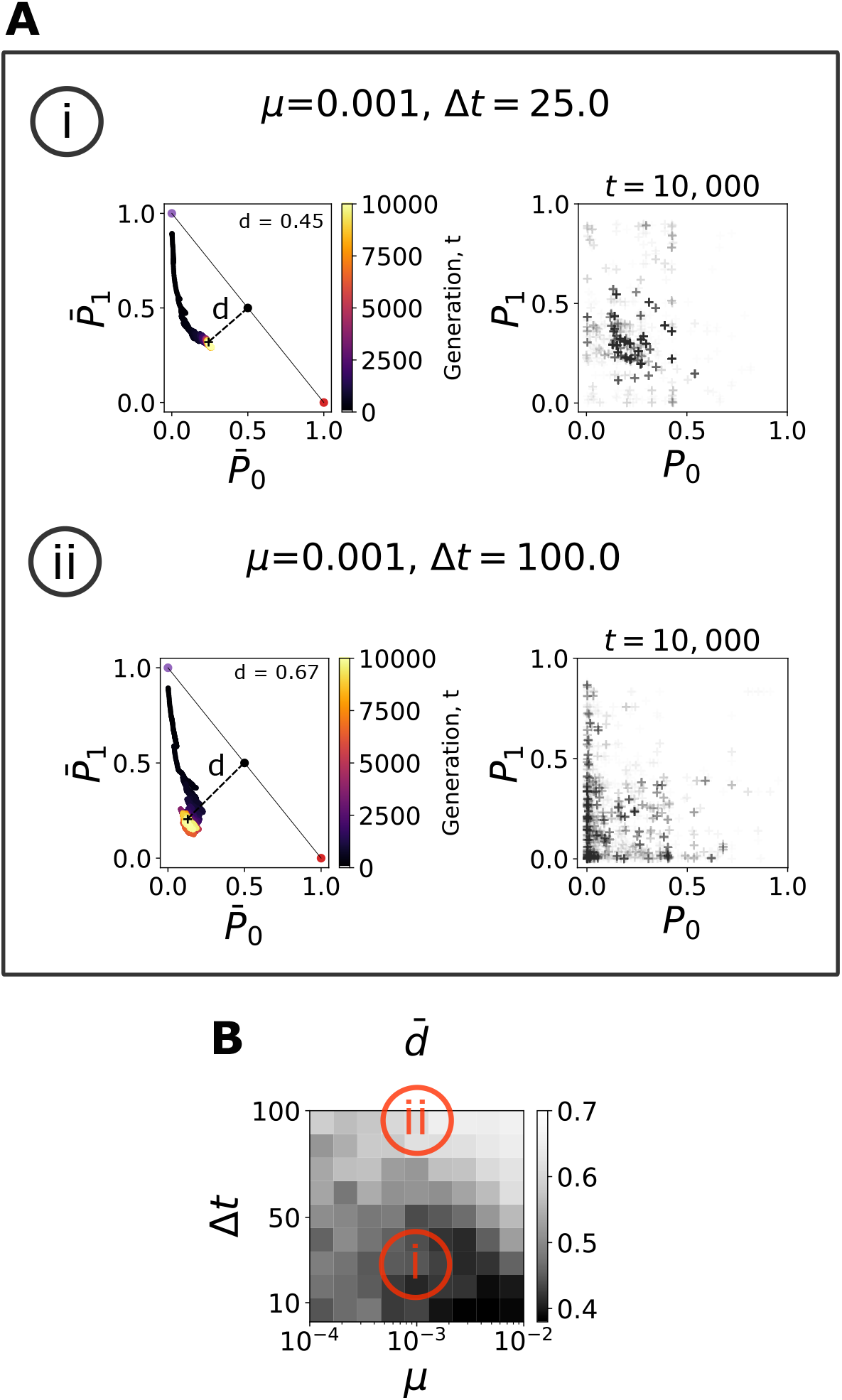
Picked examples from Fig. showing that plastic genotypes approach optimal plasticity in rapidly switching environments. **(A)** Shows the evolution over time of the degree of plasticity as measured by the mean target Boltzmann probabilities (*P̅*_0_*, P̅*_1_) of the population, for simulations (i) and (ii) in Fig.3, and the mean distance *d* at *t* = 10, 000 to the Pareto front midpoint which represents optimal plasticity. The scatter plots are accumulations of 100 simulations of the *N* = 100 population size Boltzmann probabilities at the endpoint of the simulations (*t* = 10, 000), so that the darker regions indicate higher density of points. Simulation (i) exhibits a much higher degree of plasticity as the (*P̅*_0_*, P̅*_1_) ends up much closer to the point of optimal plasticity, and the population is clustered around this end point. By contrast the final population in (ii) is much closer to the axes, indicating a low degree of plasticity and dominance of a single phenotype. **(B)** shows the parameter sweep heat map for different (*µ,* Δ*t*) of the mean distance *d̅* to optimal plasticity from the midpoint to the end of the simulation. This demonstrates a high degree of plasticity among the plastic phenotypes across the same wide range of *µ* and Δ*t* as in Fig.4 B and C. There is a strong negative correlation with the plastic abundance ratio in Fig. 4B, Pearson *r* = *−*0.63, and mean fitness in Fig. 4C, Pearson *r* = *−*0.86.

Fig. 6 shows qualitatively that a decrease in switching time, will lead towards accumulating more plasticity in populations. The measurements of plasticity are done across different target pairs for each of the 9 categories that combine ranges of Hamming distance and neutral space sizes of the RNA12 ND GP map (see Methods). In Fig. each colour represents a category, and each point is the average of all *n* = 100 points accumulated from *R_p_*(*µ,* Δ*t*), where Δ*t* is fixed. This average is calculated using the same set of 10 different *µ* values for each of the randomly selected 10 target pairs in a category, 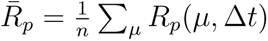. Each category of target pair shows a decrease in plasticity abundance as the switching time increases, even below the line that indicates the adapted plastic regime. Note that for Δ*t* = 1 the metric *R̅_p_* increases for most phenotype categories, except for the phenotypes with small neutral set size; the generational switching time is too fast for the average plasticity to increase, or the sample size cannot capture the behaviour well (as shown in the inset). This reflects the reduced mutational robustness and accessibility of plasticity for these phenotypes. This suggests that the emergence of plasticity may be sensitive to both the switching timescale and the underlying GP map structure, particularly for less frequent phenotypes.

**Fig 6.**
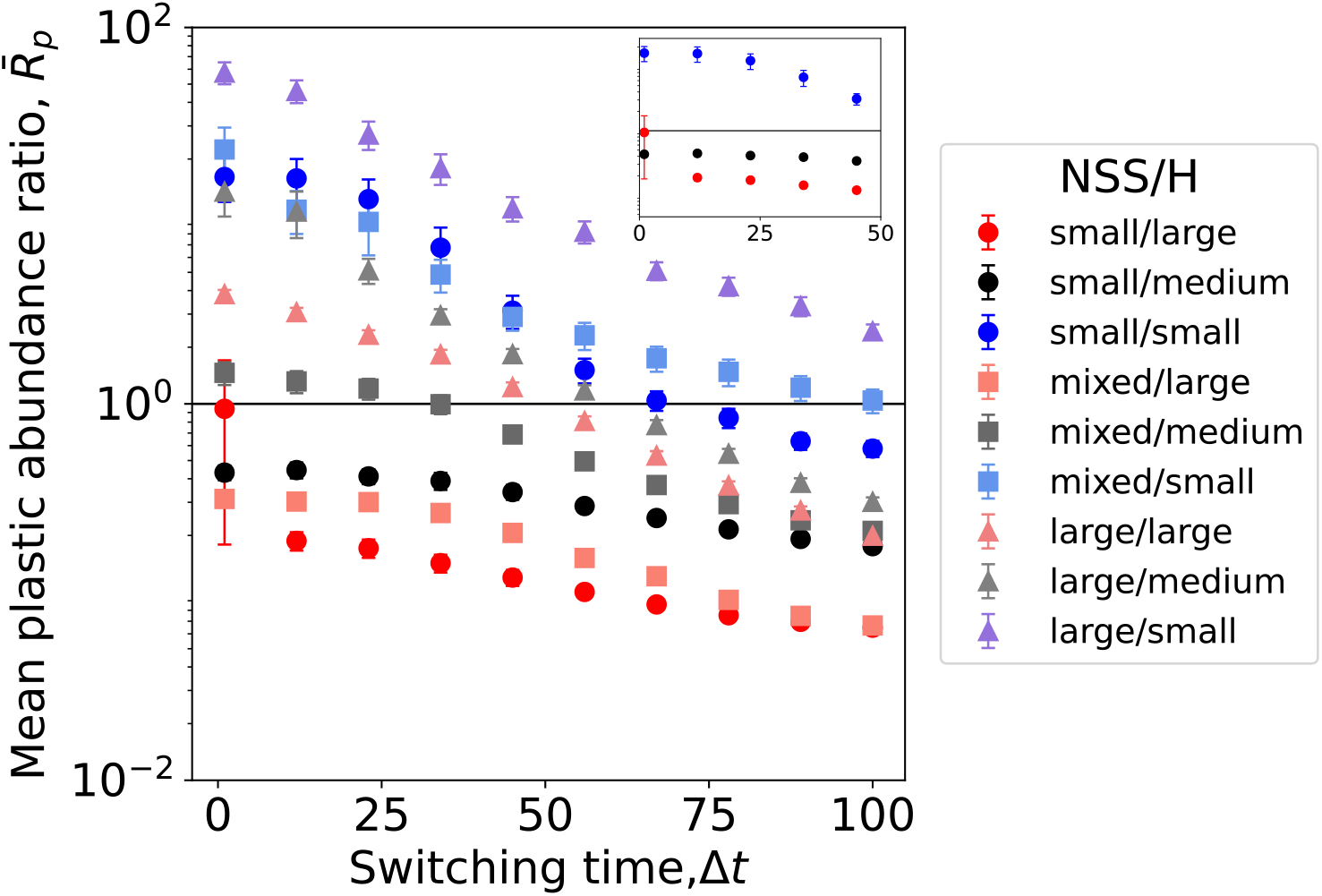
Decreasing the environmental switching time promotes the emergence of plasticity. Each point is the average of 100 points accumulated from *R_p_*(*µ,* Δ*t*), where Δ*t* is fixed (Δ*t* [1, 100]), calculated using the same set of 10 different *µ* values for each of the 10 different randomly chosen target pairs that belong to a category. The categories are defined in the Methods. The inset shows the behaviour for low switching time only for the small neutral set size phenotype pairs.

#### The structure of the genotype-phenotype map constrains the evolution of plasticity

Optimal plasticity can of course only evolve if the optimal plastic genotypes are present in the ND GP map in the first place. From the genetic correlations of the ND GP map [22, 26], we suspect that two targets’ neutral spaces are more likely to overlap if these phenotypes are common and they are similar between each other. To test this, we check that low Hamming distance and large neutral space size combine to increase plasticity by analysing the plastic abundance ratio *R_p_*for 10 different batches of 9 target pairs where each pair is randomly selected from one of the 9 different categories that combine the neutral space sizes *NSS* and Hamming distances *H* (see Methods). An example is shown in Fig.7A to show the parameter sweeps of the *R_p_* values in each target pair category. We observe an increase in *R_p_ >* 1 regions as *H* decreases and *NSS* increases, as expected. To visualise the results for the 10 different batches, the accumulated *R_p_* values for the target pairs in the 9 different categories are represented in Fig.7B. The black dotted line is *R_p_* = 1 and separates the adapted plastic (right) and non-plastic (left) *R_p_*regions. The histogram densities in Fig.7B shift towards the plastic region for large neutral spaces and small Hamming distances. Equally, we verify that the mean fitness peaks shift towards higher values with the same trend, shown in Fig.7C. This pattern is also evident in the vertical ordering of the categories observed in Fig.6.

**Fig 7.**
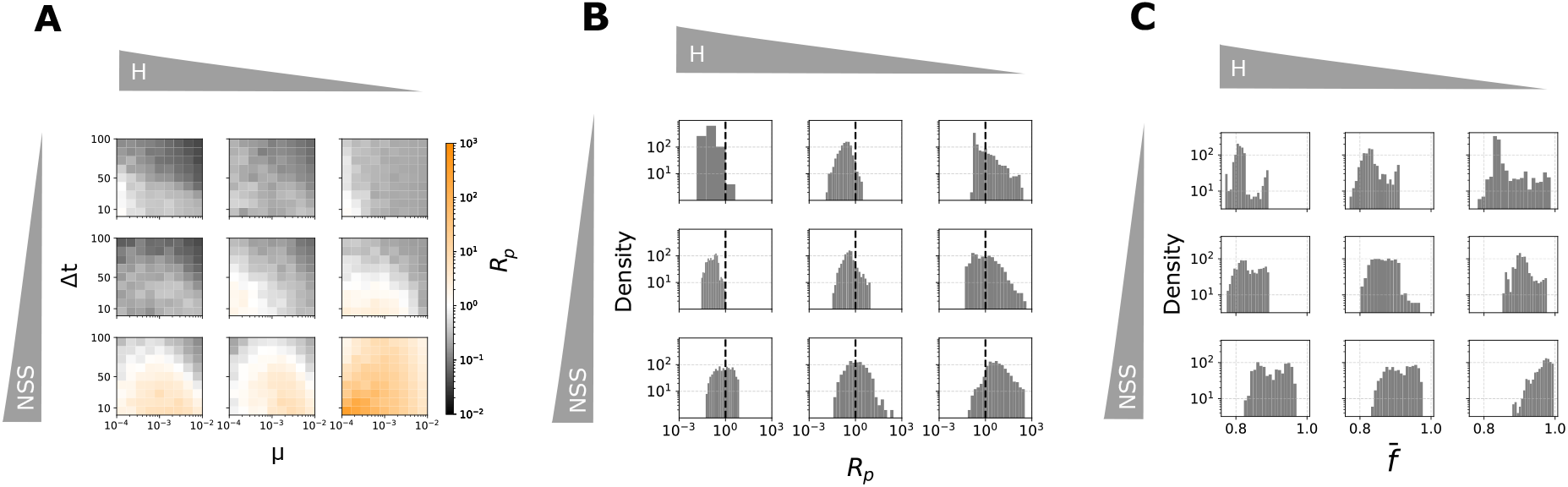
The choice of target pair has an impact in plasticity evolution determined by the ND GP map. **(A)** Heat maps of parameter sweeps (*µ,* Δ*t*) of plastic abundance ratio *R_p_*for different target pairs. Columns represent Hamming distance between secondary structures in each pair, in particular these pairs belong to *H* being large 0.8, medium 0.5, or small 0.2. The rows indicate the scale of the secondary structures’ neutral space sizes *NSS* on the RNA12 ND GP map. The top row represents small *NSS* 10^3^, middle row represents the mixed target pairs, each pair having one target secondary structure with *NSS* 10^3^ and the other with *NSS* 10^5^, and bottom row represents large *NSS* 10^5^. **(B)** Histogram of *R_p_* and mean fitness **(C)** *f̅* values of 10 different batches of 9 pairs, where each pair was randomly selected from one of the 9 categories of *NSS* and *H* that align with the categories in (A), verifying that increasing *NSS* and decreasing *H* between target pairs induces adaptive plasticity in the evolutionary dynamics of periodically switching environments.

To find the optimal plastic solution a population must navigate the ND GP map in large enough steps, i.e., with a high mutation rate, to avoid a slow navigability that could lead to not finding anything of interest, and getting trapped in local maxima (exploration catastrophe), but not so high that it exits the region of neutral space overlap (error catastrophe). The mutation rate range where the exploration is most efficient at localising the optimal plasticity will depend on the genetic correlations between the targets in the ND GP map [22, 25, 26], like previously mentioned, which can vary with structural similarity. To see an example of this, we analyse target pairs with large neutral space sizes and varying Hamming distances (see Methods), as shown in Fig. 8. Each point is the average of all *n* = 100 points accumulated from *R_p_*(*µ,* Δ*t*), where *µ* is fixed. This average is calculated using the same set of 10 different Δ*t* values for each of the randomly selected 10 target pairs in each *H* category, 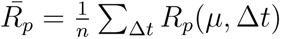. The value of *µ* that promotes higher average plasticity decreases with the Hamming distance between the targets.

**Fig 8.**
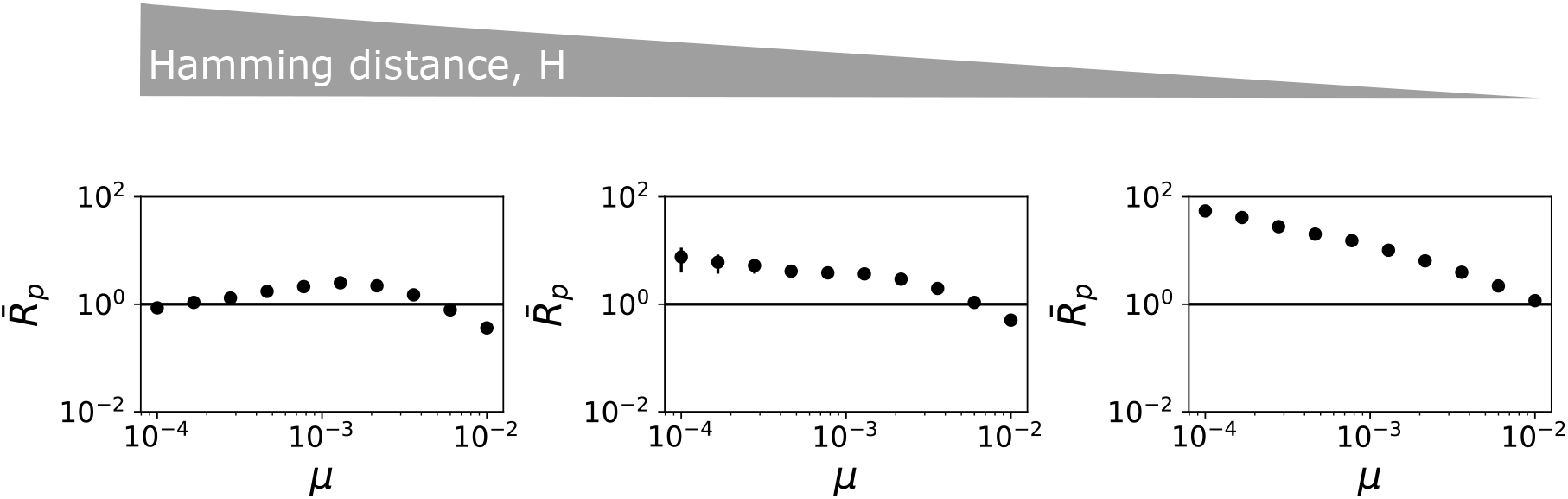
Plasticity emerges at different mutation rates depending on Hamming distance between targets. Each point is the average of 100 points accumulated from *R_p_*(*µ,* Δ*t*), where *µ* is fixed, calculated using the same set of 10 different Δ*t* values for each of the 10 different randomly chosen target pairs from each Hamming distance *H* category, in particular these pairs belong to large 0.8, medium 0.5, or small 0.2 Hamming distance. Here we are only considering targets with large neutral space sizes, *NSS ∼* 10^5^.

### Plasticity requires multiple phenotypic expressions per generation

Our fitness function picks the best option from multiple expressions of the phenotype in a given generation, through the parameter *N_p_*. This acknowledges the difference between the generational timescale and the timescale of phenotypic production. The number of samples *N_p_* reflects the selection process’s leniency: if *N_p_* = 1, only the first sampled fold determines the genotype’s fate, whereas if *N_p_ → ∞*, the sequence can access any solution within its plastic repertoire. The maximum fitness sampling function assigns the fitness *f* of a genotype by sampling *N_p_* folds *p_i_* (where *i* = 1*,…, N_p_*) from the genotype’s Boltzmann ensemble and selecting the highest Hamming fitness *f_H_*relative to a target fold *p_T_*from these samples:

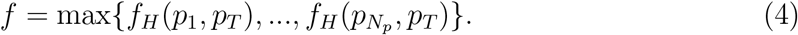

We find that plasticity only emerges if *N_p_ >* 1, but that varying it beyond that does not alter the qualitative outcomes substantially (see Fig.9), validating our choice of using a fixed value of *N_p_* = 10 for our analyses.

**Fig 9.**
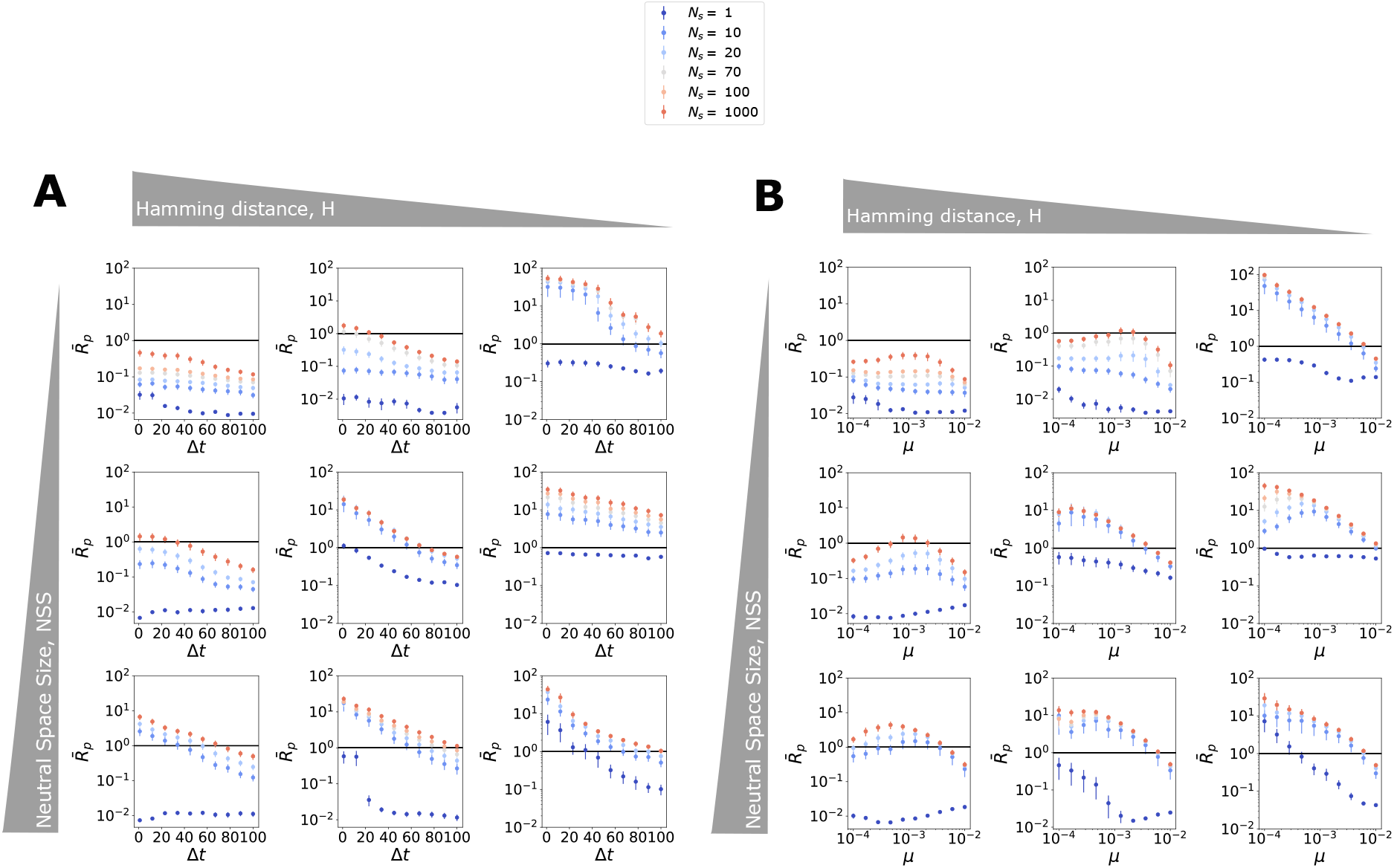
Plasticity evolves when using maximum fitness sampling function if *N_p_ >* 1. **(A)** Decrease in switching time increases average plastic abundance ratio and **(B)** for large neutral spaces, plasticity maximises at different mutation rates depending on targets’ Hamming distance. Each point is an average of 10 values of *R_p_* of 10 different (A)*µ*, (B)Δ*t* and fixed (A)Δ*t*, (B)*µ* from a randomly selected target pair of a particular category ordered by neutral set size, *NSS*, and decreasing Hamming distance, *H* (see Methods). These results align with previous results shown for *N_p_*= 10. Each colour corresponds to the *R_p_*averages for a fixed *N_p_*. The colours increase in warmth with the value of *N_p_*.

### Functional RNAs approach optimal plasticity

If naturally evolved RNAs have been selected for optimal plasticity in response to fluctuating environments, we expect that their minimum free energy (MFE) and first suboptimal structures correspond to the target phenotypes of the evolutionary process. To test this, we analyze all sequences of length *L ≤* 50 from the database fRNAdb [29], computing the Boltzmann distribution over their structural repertoire. For each sequence, we extract the MFE and first suboptimal structure along with their Boltzmann probabilities, and filter out sequences where either fold is deleterious (i.e., unfolded or non-functional). This ensures we only retain fRNAs in which both dominant structures are functionally viable.

For each retained fRNA, we compute the plasticity distance *d*, as previously defined in Eq 2, and the Hamming distance *H* between the MFE and first suboptimal structures. We further filter for high-plasticity candidates by selecting only those with *d ≤* 0.35. After this filtering, approximately 1.2% of the original fRNA database sequences remain for analysis.

To assess the statistical significance of observed plasticity, we compare each sequence’s plasticity to an empirical distribution of *d* values generated from its mutational neighborhood, sampled using a site-scanning method [30]. This method captures a representative distribution across neutral components in genotype space [30]. If statistically significant this suggests that the fRNA plasticity is unlikely to arise by random mutation alone (see Methods).

In total, 2,412 sequences of the analyzed set were found to be statistically significant (see Supplementary Data). As shown in Fig. 10, these sequences tend to exhibit higher plasticity or lower *d*, and a large proportion also display low Hamming distances *H* between MFE and suboptimal folds, consistent with predictions from our model from the RNA12 ND GP map. The plasticity of these significant sequences is not typical of general functional fRNAs as shown in Fig.10C. This confirms that the significance test successfully identifies sequences that exhibit near-optimal plasticity, whereas the full dataset contains many sequences with low plasticity.

**Fig 10.**
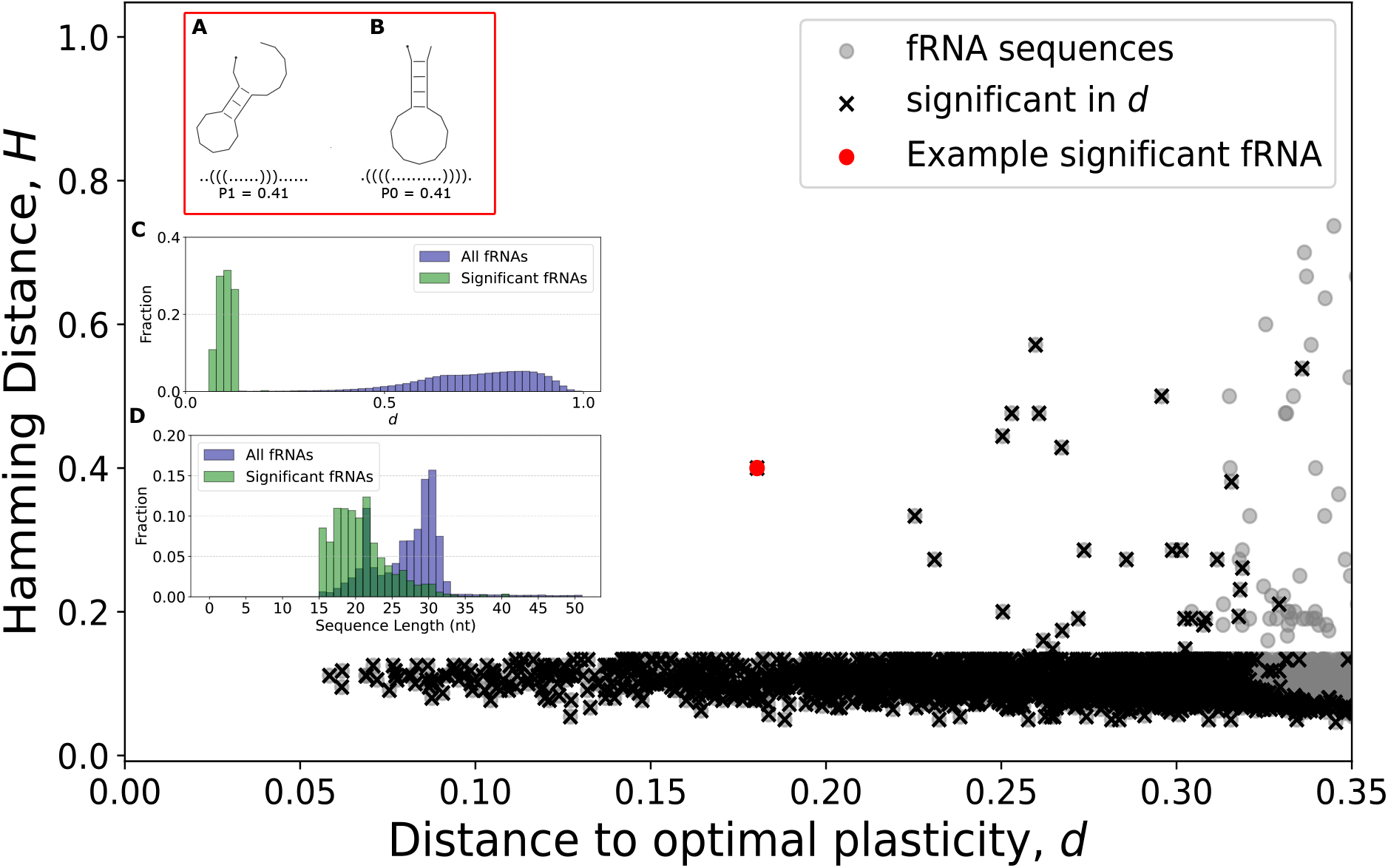
Functional RNA approaches optimal plasticity. We analysed 3,685 fRNA sequences from the fRNAdb [29] of optimal plasticity distance *d* ≤ 0.35 and sequence length *L* ≤ 50. We find 2,412 fRNA significant sequences in this set (c.f. Methods). We highlight as an example a significant sequence corresponding to Drosophila small RNA (see Supplementary Data), which corresponds to relatively low *d* and high *H*, *d* = 0.18 and *H* = 0.4, where the two ‘target’ structures are the MFE **(A)** and first suboptimal **(B)**, two different hairpin loops. **(C)** Histogram of distances *d* showing that optimal plasticity found in the significant fRNAs is not the typical plasticity found for fRNAs in general. **(D)** Histogram of sequence lengths for the significant fRNAs, showing a peak in the ranges between 18-23 nucleotides, typical of regulatory short RNAs [32, 33], compared to distribution for the full database fRNAdb [29].

An illustrative example is highlighted in Fig. 10: a short RNA from Drosophila, not yet classified functionally. The inset Fig. 10A,B shows its two dominant structures and their associated Boltzmann probabilities, yielding *d* = 0.18 and *H* = 0.4. These structures are both hairpin loops, yet clearly distinct, suggesting they may have evolved under selection pressures of alternating environments or alternating functions. While many significant fRNAs remain unclassified, including the putative conserved noncoding regions predicted by EvoFold, several have been identified as piRNAs, miRNAs, or viral regulatory elements such as those from Hepatitis C (see Supplementary data). Their plasticity may contribute to regulatory functions in response to environmental variation, such as immune pressure or developmental cues [32, 33].

The inset of Fig. 10D also shows the sequence length distribution of significant plastic fRNAs, which are predominantly 18–23 nucleotides long. These short lengths are typical of regulatory RNAs, which must function across diverse or fluctuating environments. But to respond to these alternating selection pressures, the plastic solutions must be accessible through mutation in the first place. Their short length facilitates the thermodynamic feasibility of supporting two dominant and functional structures. And, as shown in our model results, the RNA ND GP map structure allows emergence of near-optimal plasticity between two structures in fluctuating environments.

## Discussion

Phenotypic plasticity, meaning a genotype’s ability to adopt different phenotypes, is widespread in biology [34]. This phenomenon occurs at various scales and conditions, from multicellular organisms with variable traits, like leaf shape in plants, to the molecular level, where RNA and proteins change conformations. RNA, in particular, has numerous critical roles in the cell that rely on plasticity. For example, riboswitches or RNA thermometers regulate gene expression by adopting different conformations in response to ligand bindings or temperature changes, respectively [35–37]. These specific RNAs have evolved to conserve particular functions using a plastic response. In this study, we analyse the intrinsic plasticity that molecular structures have due to thermal fluctuations at the molecular scale. This thermodynamic plasticity in RNA secondary structure has been simply called in the past RNA plasticity [22]. We study this baseline plasticity under a tractable non-deterministic GP map of RNA length 12 [25]. Our research demonstrates that this RNA plasticity is adaptive in response to environmental fluctuations, depending on several factors, including the ND GP map structure. One interesting factor is that the emergence of optimal plasticity requires multiple phenotypic expressions per generation. This reflects a scenario where phenotypic switching (plasticity) is fast compared to the timescale of molecular interactions or environmental selection. This is also akin to bet-hedging [1]. Functionality, in this case, depends on the RNA being able to express the optimal structure at least some of the time, rather than on the average behavior across all structures. This winner-takes-all approach is particularly suited for regulatory and viral RNAs, where rare but structurally specific interactions can determine fitness [38, 39], which aligns with our findings of natural RNA. We observe a number of naturally evolved functional RNAs to exhibit near-optimal plasticity between two structures, unlikely to arise through mutation alone, which are short RNAs of nucleotide length range corresponding to typical regulatory and viral RNAs. Beyond natural functional RNA, our findings can be linked to research on RNA World [40] and Origin of Life, since a prevailing view is that life had to originate in fluctuating environments [41–44].

Our evolutionary model of two alternating environments may not represent a very realistic model of environmental change. Future research could investigate the evolution of RNA plasticity under unpredictable conditions [45], for different types of environmental fluctuation functions (e.g. sinusoidal), or for switching between more than two target structures. Our work therefore can be considered a baseline approach, which verifies first that plasticity is evolvable as a response to extreme change for RNA structure. This square-wave method also allows for the Pareto front framework to be implemented (see Methods).

Our current results assume an initial population already containing genotypes that have both targets (plastic genotypes) but maximizes the probability of either target structure (see Methods). It would be valuable to study the accessibility of plastic genotypes starting from a random population that lacks one of the target structures altogether. Starting from a completely random population would introduce an additional layer of complexity related to the accessibility of target phenotypes themselves, which depends strongly on GP map structure and phenotype frequency. In such a setting, the results would combine two distinct processes: (i) discovery of a plastic phenotype and (ii) emergence of high plasticity (this current work). In the target pairs we have used, the plasticity overlap is of *<* 1% (see Fig. 2), meaning the fraction of genotypes that share both structures is quite small compared to the total number of genotypes that both structures occupy, making this emergence of plasticity sensitive to dynamics and the ND GP map structure, like we see in the results. The methods for (i) on how to access the small overlap in the first place is a question we leave for future work.

However, even though our model has simplified assumptions, it contains parameters in three different relevant areas: the evolutionary dynamics, the ND GP map structure, and the selection process. It can also be applied more broadly to evolutionary systems in general, as GP maps of different biological systems exhibit similar structural properties to the RNA12 GP map analysed here [16]. This study only considers sequences of length 12. The main reason is that we wanted to do a study of plasticity evolution in an exhaustive GP map as a baseline study, and the non-deterministic GP map of RNA *L >* 12 is computationally too expensive to generate without sampling methods. The reason we want to avoid sampling is that plasticity is introduced in part of the GP map, at times, through very small probabilities. To accurately infer a genotype’s plasticity between two target pairs without generating the full Boltzmann distribution in the GP map is a study we leave for future work.

Finally, further computational work can verify the plasticity of longer length fRNA to validate our results, but ultimately we believe the next step forward could investigate the specific functional roles associated with the highly plastic fRNAs identified here. For example, one could test whether these RNAs are enriched in organisms exposed to strong environmental fluctuations, or in RNAs that function across multiple molecular contexts. Such analyses would help improve the annotation of small fRNAs, many of which remain poorly characterised, and clarify whether this near-optimal plasticity represents a general adaptive mechanism characteristic of specific classes of fRNAs with alternating functions.

## Contributions

PGG: Conceptualization, Data Curation, Formal Analysis, Software, Investigation, Methodology, Visualization, Writing – Original Draft Preparation, Writing – Review & Editing. SEA: Conceptualization, Methodology, Supervision, Writing – Review & Editing.

## Data Availability

All computational code and data is available in a public repository: https://github.com/paulagrcg/RNAplasticity(DOI: 10.5281/zenodo.13911638).

## Supplementary Information

The S1 shows the functional RNA site-scanning sample size test. The Supplementary Data with the annotations and sequences of the significant fRNAs found are shown in the file `significant sequences.xlsx` in (DOI: 10.5281/zenodo.13911638).

## Funding and Acknowledgments

PGG was supported by the “la Caixa” Foundation (Fellowship LCF/BQ/EU21/11890140) and King’s College, Cambridge. The funders had no role in study design, data collection and analysis, decision to publish, or preparation of the manuscript.

## Supplementary Information

### S1 Functional RNA site-scanning sample size

We test the stability of the empirical p-values for a set of 1000 randomly picked fRNAs by varying the number of samples used in the site-scanning procedure that constructs the null distribution. As shown in Fig.S1A, there is a substantial difference when only 100 or 1000 samples are used, compared to 10000— a standard sample size for estimating structural properties across neutral components of a GP map [1, 2]. In contrast, 5000 samples yield relatively stable p-values, as shown in Fig.S1B. We use a threshold of 0.01 for p-value stability [3, 4].

**Figure S1:**
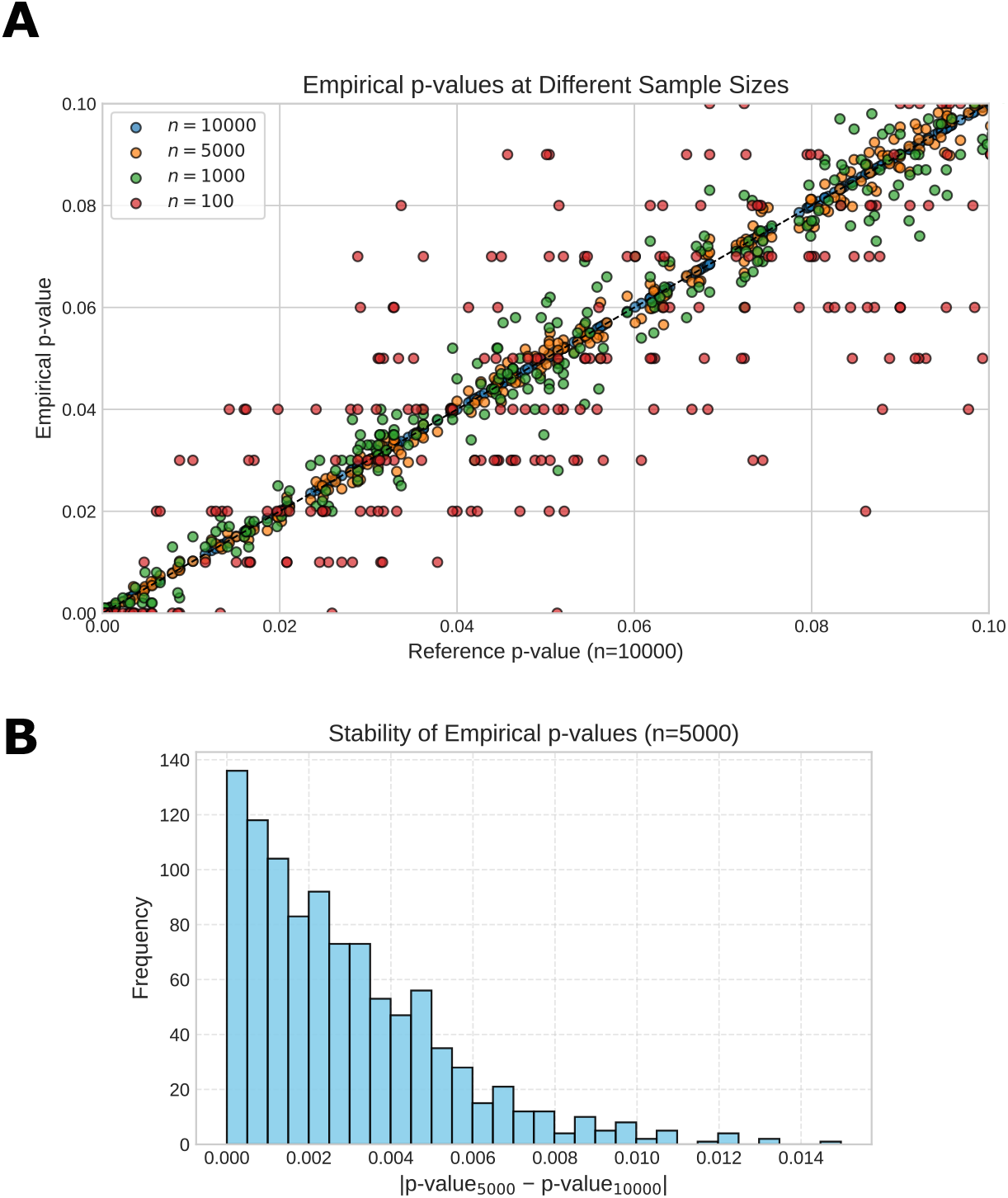
Stability of p-values for 1000 randomly picked fRNAs depending on site-scanning sample size.

## Notes

### Competing Interest Statement

The authors have declared no competing interest.

### Summary of Updates

Methods expanded: Boltzmann probability and partition function written out explicitly; choice of Hamming distance bins and population size N = 100 justified; New Figure 2 showing plastic genotypes are rare (<1% of genotypes in a target pair), with correlation r = 0.46 against genotype space size; New paragraph motivating the maximum-fitness sampling function biologically; reference added; Sample sizes for the fRNA analysis now stated (237,461 sequences of L ≤ 50; 3,685 below the cutoff; 2,412 significant); Figure 6 gains an inset for low switching time in small neutral set size pairs, with accompanying text; Figure 8 caption and surrounding text reworded; Discussion extended: RNA World and Origin of Life links, sinusoidal fluctuation as a future direction, and new paragraphs on the initial population assumption, the restriction to L = 12, and future functional characterisation of the plastic fRNAs; Contributions and Funding sections added; references updated.

## References

1. L. A. Meyers and J. J. Bull, “Fighting change with change: adaptive variation in an uncertain world”, Trends in Ecology & Evolution 17, 551–557 (2002).

2. R. Levins, Evolution in Changing Environments: Some Theoretical Explorations. (MPB- 2) (Princeton University Press, 1968).

3. E. Kussell and S. Leibler, “Phenotypic Diversity, Population Growth, and Information in Fluctuating Environments”, Science 309, 2075–2078 (2005).

4. O. Shoval, H. Sheftel, G. Shinar, Y. Hart, O. Ramote, A. Mayo, E. Dekel, K. Kavanagh, and U. Alon, “Evolutionary Trade-Offs, Pareto Optimality, and the Geometry of Phenotype Space”, Science 336, 1157–1160 (2012).

5. M. Tikhonov, S. Kachru, and D. S. Fisher, “A model for the interplay between plastic tradeoffs and evolution in changing environments”, Proceedings of the National Academy of Sciences 117, 8934–8940 (2020).

6. L. W. Ancel, “Undermining the Baldwin Expediting Effect: Does Phenotypic Plasticity Accelerate Evolution?”, Theoretical Population Biology 58, 307–319 (2000).

7. R. Lande, “Adaptation to an extraordinary environment by evolution of phenotypic plasticity and genetic assimilation”, Journal of Evolutionary Biology 22, 1435–1446 (2009).

8. M. Pigliucci, “Evolution of phenotypic plasticity: where are we going now?”,Ecology & Evolution 20, 481–486 (2005).

9. S. Manrubia, J. A. Cuesta, J. Aguirre, S. E. Ahnert, L. Altenberg, A. V. Cano, P. Catalán, R. Diaz-Uriarte, S. F. Elena, J. A. García-Martín, P. Hogeweg, B. S. Khatri, J. Krug, A. A. Louis, N. S. Martin, J. L. Payne, M. J. Tarnowski, and M. Weiß, “From genotypes to organisms: State-of-the-art and perspectives of a cornerstone in evolutionary dynamics”, Physics of Life Reviews 38, 55–106 (2021).

10. K. Dingle, F. Ghaddar, P. Šulc, and A. A. Louis, “Phenotype Bias Determines How Natural RNA Structures Occupy the Morphospace of All Possible Shapes”, Biology and Evolution 39, 10.1093/molbev/msab280 (2021).

11. P. Schuster, W. Fontana, P. F. Stadler, and I. L. Hofacker, “From sequences to shapes and back: a case study in RNA secondary structures”, Proceedings of the Royal Society of London. Series B: Biological Sciences 255, 279–284 (1994).

12. P. Schuster, “Prediction of RNA secondary structures: from theory to models and real molecules”, Reports on Progress in Physics 69, 1419–1477 (2006).

13. A. Wagner, “Robustness and evolvability: a paradox resolved”,Society B: Biological Sciences 275, 91–100 (2008).

14. S. Schaper and A. A. Louis, “The Arrival of the Frequent: How Bias in Genotype-Phenotype Maps Can Steer Populations to Local Optima”, PLoS ONE 9, e86635 (2014).

15. S. F. Greenbury, S. Schaper, S. E. Ahnert, and A. A. Louis, “Genetic Correlations Greatly Increase Mutational Robustness and Can Both Reduce and Enhance Evolvability”, PLOS Computational Biology 12, edited by R. A. Goldstein, e1004773 (2016).

16. S. E. Ahnert, “Structural properties of genotype–phenotype maps”,Royal Society Interface 14, 20170275 (2017).

17. R. Lorenz, S. H. Bernhart, C. H. z. Siederdissen, H. Tafer, C. Flamm, P. F. Stadler, and I. L. Hofacker, “ViennaRNA Package 2.0”,Algorithms for Molecular Biology 6, 26 (2011).

18. H. Schwalbe, J. Buck, B. Fürtig, J. Noeske, and J. Wöhnert, “Structures of RNA Switches: Insight into Molecular Recognition and Tertiary Structure”, Angewandte Chemie International Edition 46, 1212–1219 (2007).

19. M. Doetsch, R. Schroeder, and B. Fürtig, “Transient RNA–protein interactions in RNA folding”, The FEBS Journal 278, 1634–1642 (2011).

20. S. F. Greenbury, A. A. Louis, and S. E. Ahnert, “The structure of genotype-phenotype maps makes fitness landscapes navigable”, Nature Ecology & Evolution, 1–11 (2022).

21. I. G. Johnston, K. Dingle, S. F. Greenbury, C. Q. Camargo, J. P. K. Doye, S. E. Ahnert, and A. A. Louis, “Symmetry and simplicity spontaneously emerge from the algorithmic nature of evolution.”,

22. L. W. Ancel and W. Fontana, “Plasticity, evolvability, and modularity in RNA”,of Experimental Zoology 288, 242–283 (2000).

23. N. Vaidya and N. Lehman, “One RNA plays three roles to provide catalytic activity to a group I intron lacking an endogenous internal guide sequence”,

24. E. A. Schultes and D. P. Bartel, “One Sequence, Two Ribozymes: Implications for the Emergence of New Ribozyme Folds”, Science 289, 448–452 (2000).

25. P. García-Galindo, S. E. Ahnert, and N. S. Martin, “The non-deterministic genotype–phenotype map of RNA secondary structure”,

26. S. Wuchty, W. Fontana, I. L. Hofacker, and P. Schuster, “Complete suboptimal folding of RNA and the stability of secondary structures”, Biopolymers 49, 145–165 (1999).

27. W. J. Ewens, Mathematical Population Genetics, edited by S. S. Antman, J. E. Marsden, L. Sirovich, and S. Wiggins, Vol. 27, Interdisciplinary Applied Mathematics (Springer New York, New York, NY, 2004).

28. Q. Zhang, A. C. Stelzer, C. K. Fisher, and H. M. Al-Hashimi, “Visualizing spatially correlated dynamics that directs RNA conformational transitions”,

29. T. Kin, K. Yamada, G. Terai, H. Okida, Y. Yoshinari, Y. Ono, A. Kojima, Y. Kimura, T. Komori, and K. Asai, “fRNAdb: a platform for mining/annotating functional RNA candidates from non-coding RNA sequences”,

30. M. Weiß and S. E. Ahnert, “Using small samples to estimate neutral component size and robustness in the genotype–phenotype map of RNA secondary structure”, en, of The Royal Society Interface 17, 20190784 (2020).

31. Y. Benjamini and Y. Hochberg, “Controlling the False Discovery Rate: A Practical and Powerful Approach to Multiple Testing”,Journal of the Royal Statistical Society: Series B (Methodological) 57, 289–300 (1995).

32. D. P. Bartel, “MicroRNAs: Target Recognition and Regulatory Functions”, Cell 136, 215–233 (2009).

33. D. M. Ozata, I. Gainetdinov, A. Zoch, D. O’Carroll, and P. D. Zamore, “PIWI-interacting RNAs: small RNAs with big functions”, en, Nature Reviews Genetics 20, 89–108 (2019).

34. M. J. West-Eberhard, Developmental Plasticity and Evolution (Oxford University Press, Mar. 2003).

35. M. Mandal and R. R. Breaker, “Gene regulation by riboswitches”, Nature Reviews Molecular Cell Biology 5, 451–463 (2004).

36. J. Kortmann and F. Narberhaus, “Bacterial RNA thermometers: molecular zippers and switches”, Nature Reviews Microbiology 10, 255–265 (2012).

37. S. E. Thomas, M. Balcerowicz, and B. Y.-W. Chung, “RNA structure mediated thermoregulation: What can we learn from plants?”, Nature Reviews Microbiology 10, 255–265 (2012).

38. A. M. Mustoe, S. Busan, G. M. Rice, C. E. Hajdin, B. K. Peterson, V. M. Ruda, N. Kubica, R. Nutiu, J. L. Baryza, and K. M. Weeks, “Pervasive Regulatory Functions of mRNA Structure Revealed by High-Resolution SHAPE Probing”, Cell 173, 181–195.e18 (2018).

39. S. Richter, H. Cao, and T. M. Rana, “Specific HIV-1 TAR RNA Loop Sequence and Functional Groups Are Required for Human Cyclin T1TatTAR Ternary Complex Formation”, Biochemistry 41, 6391–6397 (2002).

40. Á. Kun, A. Szilágyi, B. Könnyű G. Boza, I. Zachar, and E. Szathmáry, “The dynamics of the RNA world: insights and challenges”, en, Annals of the New York Academy of Sciences 1341, 75–95 (2015).

41. P. Higgs, “The Effect of Limited Diffusion and Wet–Dry Cycling on Reversible Polymerization Reactions: Implications for Prebiotic Synthesis of Nucleic Acids”, en, Life 6, 24 (2016).

42. S. Roy, N. V. Bapat, J. Derr, S. Rajamani, and S. Sengupta, “Emergence of ribozyme and tRNA-like structures from mineral-rich muddy pools on prebiotic earth”, en, Journal of Theoretical Biology 506, 110446 (2020).

43. A. V. Tkachenko and S. Maslov, “Spontaneous emergence of autocatalytic information-coding polymers”, en, The Journal of Chemical Physics 143, 045102 (2015).

44. C. Alejandre, A. Aguirre-Tamaral, C. Briones, and J. Aguirre, “Polymerization and replication of primordial RNA induced by clay-water interface dynamics”, en, Communications Chemistry 8, 236 (2025).

45. B. Xue, P. Sartori, and S. Leibler, “Environment-to-phenotype mapping and adaptation strategies in varying environments”, Proceedings of the National Academy of Sciences 116, 13847–13855 (2019)

## References

1. P. García-Galindo, S. E. Ahnert, and N. S. Martin, “The non-deterministic genotype–phenotype map of RNA secondary structure”, Journal of The Royal Society Interface 20, 20230132 (2023).

2. M. Weiß and S. E. Ahnert, “Using small samples to estimate neutral component size and robustness in the genotype–phenotype map of RNA secondary structure”, en, Journal of The Royal Society Interface 17, 20190784 (2020).

3. B. Phipson and G. K. Smyth, “Permutation P-values Should Never Be Zero: Calculating Exact P-values When Permutations Are Randomly Drawn”, en, Statistical Applications in Genetics and Molecular Biology 9, Publisher: De Gruyter, 10.2202/1544-6115.1585 (2010).

4. B. V. North, D. Curtis, and P. C. Sham, “A Note on the Calculation of Empirical *P* Values from Monte Carlo Procedures”, The American Journal of Human Genetics 71, 439–441 (2002).

